# Long-term passaging of replication competent pseudo-typed SARS-CoV-2 reveals the antiviral breadth of monoclonal and bispecific antibody cocktails

**DOI:** 10.1101/2022.10.05.511057

**Authors:** Hang Ma, Huifang Zong, Junjun Liu, Yali Yue, Yong Ke, Yunji Liao, Haoneng Tang, Lei Wang, Shusheng Wang, Yunsheng Yuan, Mingyuan Wu, Yanlin Bian, Baohong Zhang, Haiyang Yin, Hua Jiang, Tao Sun, Lei Han, Yueqing Xie, Jianwei Zhu

## Abstract

The continuous emergence of severe acute respiratory syndrome coronavirus 2 (SARS-CoV-2) variants poses challenges to the effectiveness of neutralizing antibodies. Rational design of antibody cocktails is a realizable approach addressing viral immune evasion. However, evaluating the breadth of antibody cocktails is essential for understanding the development potential. Here, based on a replication competent vesicular stomatitis virus model that incorporates the spike of SARS-CoV-2 (VSV-SARS-CoV-2), we evaluated the breadth of a number of antibody cocktails consisting of monoclonal antibodies and bispecific antibodies by long-term passaging the virus in the presence of the cocktails. Results from over two-month passaging of the virus showed that 9E12+10D4+2G1 and 7B9-9D11+2G1 from these cocktails were highly resistant to random mutation, and there was no breakthrough after 30 rounds of passaging. As a control, antibody REGN10933 was broken through in the third passage. Next generation sequencing was performed and several critical mutations related to viral evasion were identified. These mutations caused a decrease in neutralization efficiency, but the reduced replication rate and ACE2 susceptibility of the mutant virus suggested that they might not have the potential to become epidemic strains. The 9E12+10D4+2G1 and 7B9-9D11+2G1 cocktails that picked from the VSV-SARS-CoV-2 system efficiently neutralized all current variants of concern and variants of interest including the most recent variants Delta and Omicron, as well as SARS-CoV-1. Our results highlight the feasibility of using the VSV-SARS-CoV-2 system to develop SARS-CoV-2 antibody cocktails and provide a reference for the clinical selection of therapeutic strategies to address the mutational escape of SARS-CoV-2.

## Introduction

Coronavirus disease 2019 (COVID-19) is caused by the infection of severe acute respiratory syndrome coronavirus 2 (SARS-CoV-2) and has been a persistent and large-scale pandemic around the world, causing hundreds of millions of infections and millions of deaths^1–3^. SARS-CoV-2 encodes an exoribonuclease in nonstructural protein 14 (nsp14-ExoN) that is proposed to have the function of nucleotide proofreading^4^. However, the virus is still in fast mutation. The World Health Organization (WHO) labeled several SARS-CoV-2 variants as variant of concern (VOC) and variant of interest (VOI) based on their potential transmissibility and risk. At present, there are five VOCs, B.1.1.7 (Alpha), B.1.351 (Beta), P.1 (Gamma), B.1.617.2 (Delta) and B. 1.1.529 (Omicron), respectively^5,6^. These variants contain mutations that contribute to the profile alteration of the virus such as L452R, S477N, T478K, E484K and N501Y. The L452R stabilized the conformation of the S protein^7^ and led to an increased affinity between the virus and angiotensin converting enzyme 2 (ACE2)^8^. Substitution in S477 with asparagine (N) resulted in an additional glycosylation of receptor binding domain (RBD)^9^. The basic charged lysine (K) substitution in T478 is thought to increase the electrostatic potential of spike (S) protein^10^. These mutations, individually or in combination, are high likely to give rise to the immune escape of virus from vaccines and neutralizing antibodies^11,12^.

Abundant neutralizing antibodies targeting conserved epitopes of the S protein have been identified in the recent two years^13,14^, but mutations at key sites may still lead to the loss of activity. The emergence of Omicron variant with numerous and scattered mutations in the N-terminal domain (NTD) and RBD renders no effective to the existing neutralizing antibodies^15,16^. Researchers compared the activity of antibodies in clinic against the Omicron and found that most of them lost neutralization activity except Sotrovimab (VIR-7831) and DXP-604 retained partial activity^17^. Another study came to a similar conclusion that only Brii-198 had better neutralization against Omicron compared to D614G virus^18^, which raised concerns about the feasibility of using a single neutralizing antibody to address the mutation of SARS-CoV-2.

The combinational use of antibodies targeting different epitopes is a widely accepted strategy. Antibodies in different epitopes can bind to S protein simultaneously, thus the virus is more difficult to break through. Besides, antibodies in different epitopes may have diverse mechanisms. The combination of CoV2-06 and CoV2-14 was effective in preventing infection by SARS-CoV-2 variants, which was far superior to the two antibodies alone^19^. The combination of REGN10987 and REGN10933 protected hamsters and non-human primates prophylactically and therapeutically, and effectively reduced viral load and pathological changes^20^. However, combinations by less breadth antibodies still present an escape risk. Besides, antibody combinations designed based on current variants always exhibit broad neutralizing activity, but their breadth is difficult to fully understand and does not provide guidance for future variants.

In this study, we formulated several sets of antibody cocktails from our SARS-CoV-2 antibody repertoire and evaluated the viral escape ability from these cocktails in a replication-competent vesicular stomatitis virus model with S protein replacement (VSV-SARS-CoV-2) through long-term passaging.^21^ Considering the combination of two or more antibodies may potentially complicate the product components, introduce complex manufacturing procedures, and increase production costs, we developed some of these monoclonal antibodies into a bunch of bispecific antibodies using our BAPTS bispecific antibodies development platform^22,23^.

## Results

### Selection of antibodies for combinational use

In our previous study^21^, we identified a broad-spectrum neutralizing antibody 2G1 that targets the tip of RBD through a small contact surface from our SARS-CoV-2 antibody repertoire, which makes it less possible in epitope clash with other antibodies and thus suitable for combinational use. We performed competition ELISA to map the epitope information of antibodies. As expected, all of the tested antibodies competed with themselves (Figure 1a). Antibodies in the repertoire could be roughly divided into three groups according to their competition relation, of which antibody 2G1 was in the group 3. Then, we examined whether these antibodies could cross bind to SARS-CoV-1 as the ability the binding to the *Sarbecovirus* could be a reflection of conserved activity by an anti-SARS-CoV-2 antibody. As shown in Figure 1b, all antibodies in group 1 and 2, but not in group 3, efficiently bound to SARS-CoV-1 S protein at the concentration of 0.1 μg/mL. We then performed pseudo-typed SARS-CoV-1 virus neutralization (Figure 1c). Antibodies 7G10, 8G4, 9E12, 9D11, 8C12 and 9A6 showed high neutralizing activity to SARS-CoV-1, while most of them were in epitope group 1. Subsequently, we explored the ACE2 competition profile of these antibodies and found that antibodies in group 1 and 3 showed excellent ACE2 competition while the group 2 antibodies had little or negligeable level of the competition(Figure 1d).

**Figure 1.**
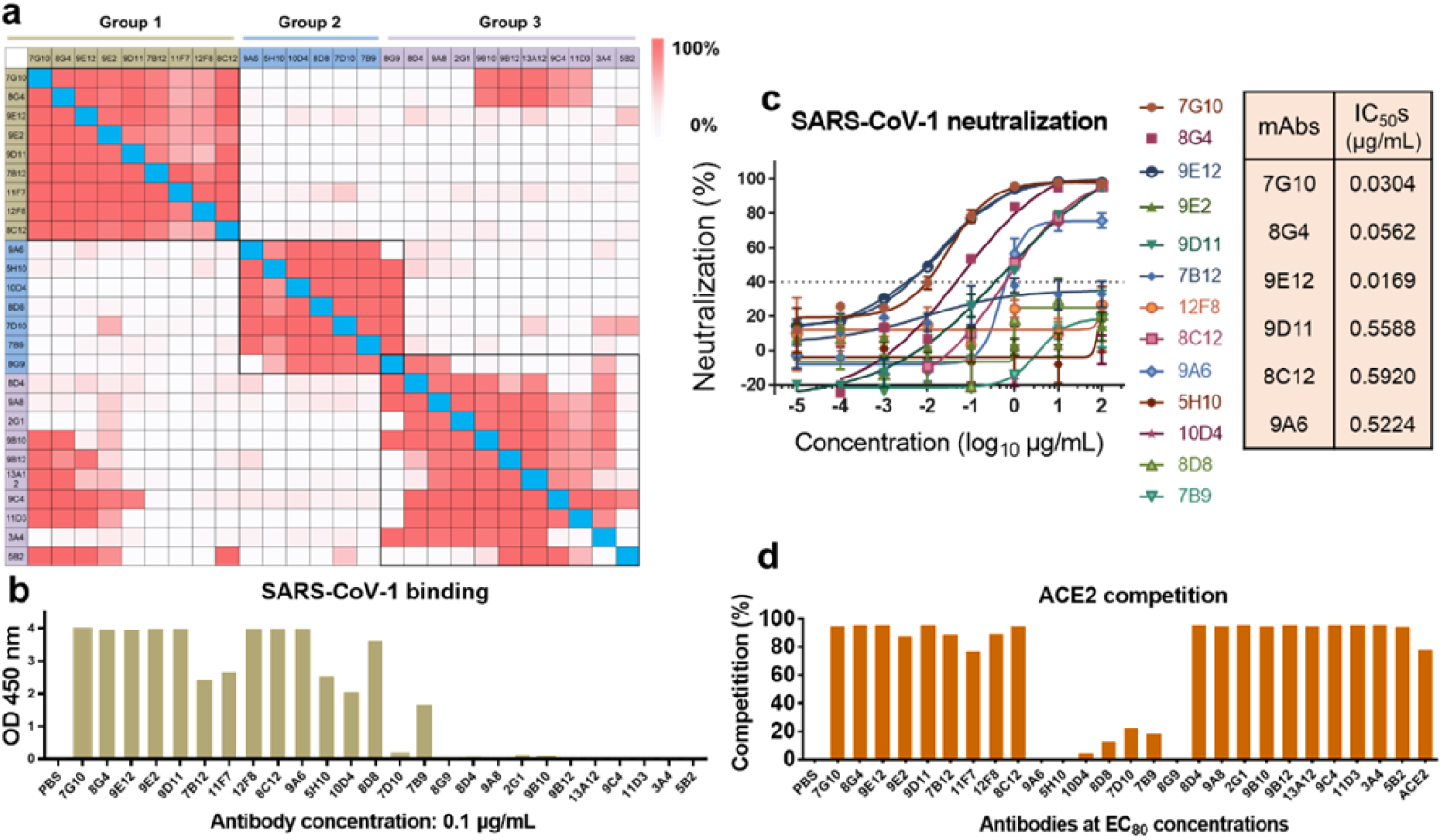
Selection of antibody cocktails from the SARS-CoV-2 neutralizing antibody repertoire. **a,** Epitope mapping of the antibody repertoire using a competition ELISA. A heatmap was used to show the competition percentages between two antibodies. The epitopes of antibodies was roughly classified into three groups according to their competitive relations. **b,** Binding to SARS-CoV-1 S protein by antibodies in the repertoire. Trimeric SARS-CoV-1 S protein was coated on 96-well ELISA plates and incubated with 0.1 μg/mL of neutralizing antibodies. The binding was detected using a HRP-labeled goat anti-human IgG (Fc specific) antibody. **c,** Concentration-dependent neutralization of SARS-CoV-1 pseudovirus by neutralizing antibodies. Serial diluted antibodies were incubated with the virus and used to infect 293T-ACE2 cells. The infection was quantified using a fluorescence quantification kit. Data from two replicates are shown as mean ± S.D. **d,** Antibodies competitively blocked the ACE2 binding to SARS-CoV-2 S trimer as measured by ELISA. Recombinant human ACE2 protein and phosphate buffer solution (PBS) were used as controls.

These results revealed certain regularities. Antibodies in group 1 were able to bind to SARS-CoV-1 spike and neutralized the virus by blocking ACE2 interaction. The group 2 antibodies targeted non-ACE2 epitope in the spike of SARS-CoV-1 but were incapable of neutralizing the virus. The group 3 antibodies only showed SARS-CoV-2 binding and neutralizing ability, instead of SARS-CoV-1. To minimize the possibility of epitope clash, we intended to select antibodies from the group 1and 2 to make up combinations with 2G1. Finally, antibodies 9E12, 9D11, 8G4 and 11F7 from group 1 and antibodies 10D4 and 7B9 from group 2 were selected. Then, we used two formulating strategies, one was two monoclonal antibodies paired with 2G1, another was a bispecific antibody paired with 2G1. We designed bispecific antibodies 7B9-9D11, 10D4-8G4 and 10D4-11F7 using the BAPTS platform^22^. Eventually, four antibody cocktails were generated, including 9E12+10D4+2G1, 7B9-9D11+2G1, 10D4-8G4+2G1 and 10D4-11F7+2G1.

### Testing the tolerance of antibody cocktails by long-term virus passaging

We then used a VSV-SARS-CoV-2 system that can generate random mutations during replication to investigate the antimutagenic potential of the four cocktails (Figure 2a). The virus in this investigation experienced over 30 passages, and the entire 30 passaging cycles costed more than 60 days, which offered the virus an opportunity generating sufficient mutations in adequate time. The VSV-SARS-CoV-2 broke through the protection of monotherapy by 9E12, 10D4 and 2G1 at the 9^th^, 1^st^ and 4^th^ passages, respectively (Figure 2b). The emergency use authorization (EUA) antibody REGN10933, which has a similar epitope with 2G1, was used as a control^21^. The virus broken through REGN10933 at the 3^rd^ passage. As for the three bispecific antibodies, 10D4-8G4 lost activity at the 4^th^ passage, 10D4-11F7 lost neutralization at the 1^st^ passage. By contrast, 7B9-9D11 showed outstanding tolerance and was not broken through even at the designed endpoint of the 30^th^ passage, despite the reduced neutralization.

**Figure 2.**
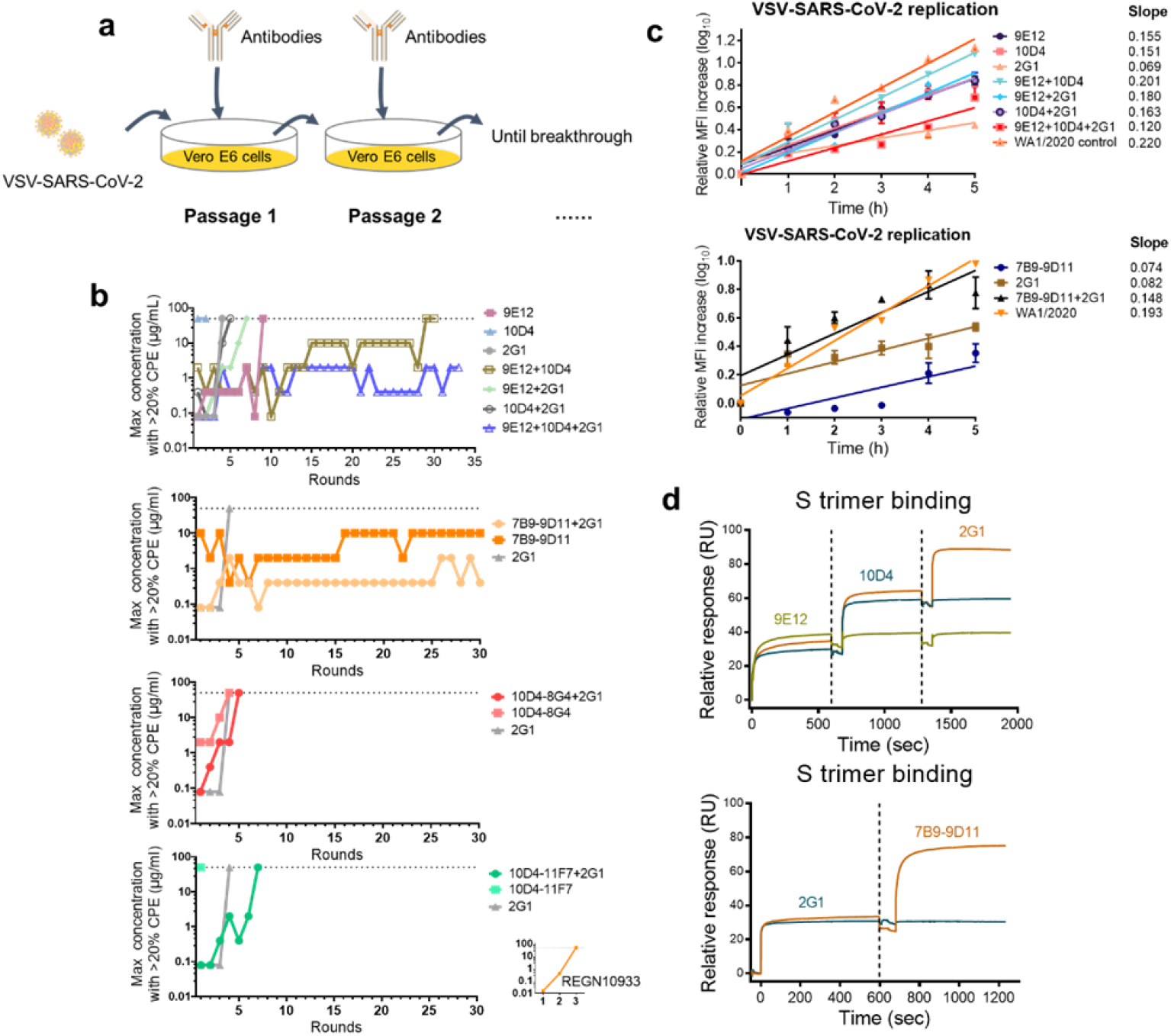
Long-term monitoring of the antibody cocktails protect cells from the infection of vesicular stomatitis viruses with S protein replacement (VSV-SARS-CoV-2). **a**, The passaging scheme of the long-term protection of antibody cocktails in a Vero E6 cell model. **b,** Antibody cocktails were 5-fold serially diluted from 100 μg/mL and mixed with VSV-SARS-CoV-2 before infecting Vero E6 cells. After 48 h, the cytopathic effect in each well was examined and medium from the highest concentration wells that the viruses showed positive infection was collected for the next round of passage. An authorized antibody REGN10933 was used as control. **c,** Comparison of the replication rates of VSV-SARS-CoV-2 virus that broke through the protection of antibody cocktail. The replication rates were calculated according to the mean fluorescence intensity measured by flow cytometry. **d,** Simultaneous binding of antibodies to WA1/2020 S trimer as measured by surface plasmon resonance. The simultaneous binding was confirmed by the accumulation of response values from different rounds of flow.

Combinational therapy by 9E12+10D4 significantly enhanced the anti-mutation ability and prevented the infection of the virus until the 29^th^ passage (Figure 2b). The triple combination by 9E12+10D4+2G1 showed more stronger resistance to viral mutation and had not been broken through even at the 33^rd^ passage. The 7B9-9D11+2G1 cocktail showed not only outstanding tolerance, but also good neutralizing activity. The results indicated that the antibody cocktails we designed improved tolerance to virus from monotherapy substantially. However, not all combinations were promising in dealing with mutational escape. Combinations by 10D4-8G4+2G1 and 10D4-11F7+2G1 increased resistance to mutations merely, and were broke through at 5^th^ and 7^th^, respectively.

The replication rate of a virus is in correlation with its transmissibility. In view of the good tolerance of combinations by 9E12+10D4+2G1 and 7B9-9D11+2G1, we collected the passaged viruses and measured their replication rates (Figure 2c). The passaged viruses generally had lower growth rates than the WA1/2020 control. The slope of the replication curve of viruses from 9E12+10D4+2G1 and 7B9-9D11+2G1 were 0.120 (versus 0.220 in WA1/2020 control) and 0.148 (versus 0.193 in WA1/2020 control). Notably, although the virus broke through the protection of monotherapy by 2G1 quickly, the replication rate of the mutant viruse was relatively slower (slop = 0.069 and 0.082). The slowed replication rates of these viruses might suggest less possibility of pandemic in the real world.

Subsequently, we verified whether antibodies in 9E12+10D4+2G1 and 7B9-9D11+2G1 cocktails could bind to S protein simultaneously using surface plasmon resonance (SPR). When 9E12, 10D4 and 2G1, or 7B9-9D11 and 2G1 flowed through the antigen-coated chip sequentially, the response signal increased accordingly, while that in the control channels remained unchanged, indicating that antibodies in both of the two cocktails could bind to S protein simultaneously (Figure 2d).

### ACE2 susceptibility of passaged VSV-SARS-CoV-2 viruses

The binding of S protein to ACE2 on the cell surface is a key step of virus invasion. Higher the affinity of S protein to ACE2 means the easier for the virus infecting cells. Meanwhile, the higher ACE2 affinity makes the virus easier to be neutralized by recombinant ACE2 protein in the medium. Thus, the susceptibility of viruses to recombinant ACE2 protein can be used to evaluate the difficulty of viruses invading cells. To further verify the reliability of the anti-mutational potential of the VSV-SARS-CoV-2 system in evaluating antibody cocktails, we examined the ACE2 susceptibility for 9E12+10D4+2G1 and 7B9-9D11+2G1. As shown in Figure 3a and b, none of the mutant viruses had increased ACE2 susceptibility compared with the WA1/2020 control, indicating that mutations in these viruses did not increase the affinity to ACE2 and may not lead to an increased risk of transmission. Specifically, viruses from pressurized passaging by 9E12+10D4, 7B9-9D11 and 7B9-9D11+2G1 showed more than 9-fold decreases in ACE2 sensitivity, and the 9E12+10D4 virus decreased by more than 27-fold. The reduced ACE2 susceptibility is correlated with the slowed cell invasion; this should be a contributive factor in the reduction of virus replication rates that observed in Figure 2c. Reduced ACE2 susceptibility may result in degraded viral transmissibility without presenting an epidemic risk. Therefore, even though the mutant viruses escaped or weakened the protection of the two antibody cocktails, transmission of these viruses may not occur in nature due to the reduced transmissibility.

**Figure 3.**
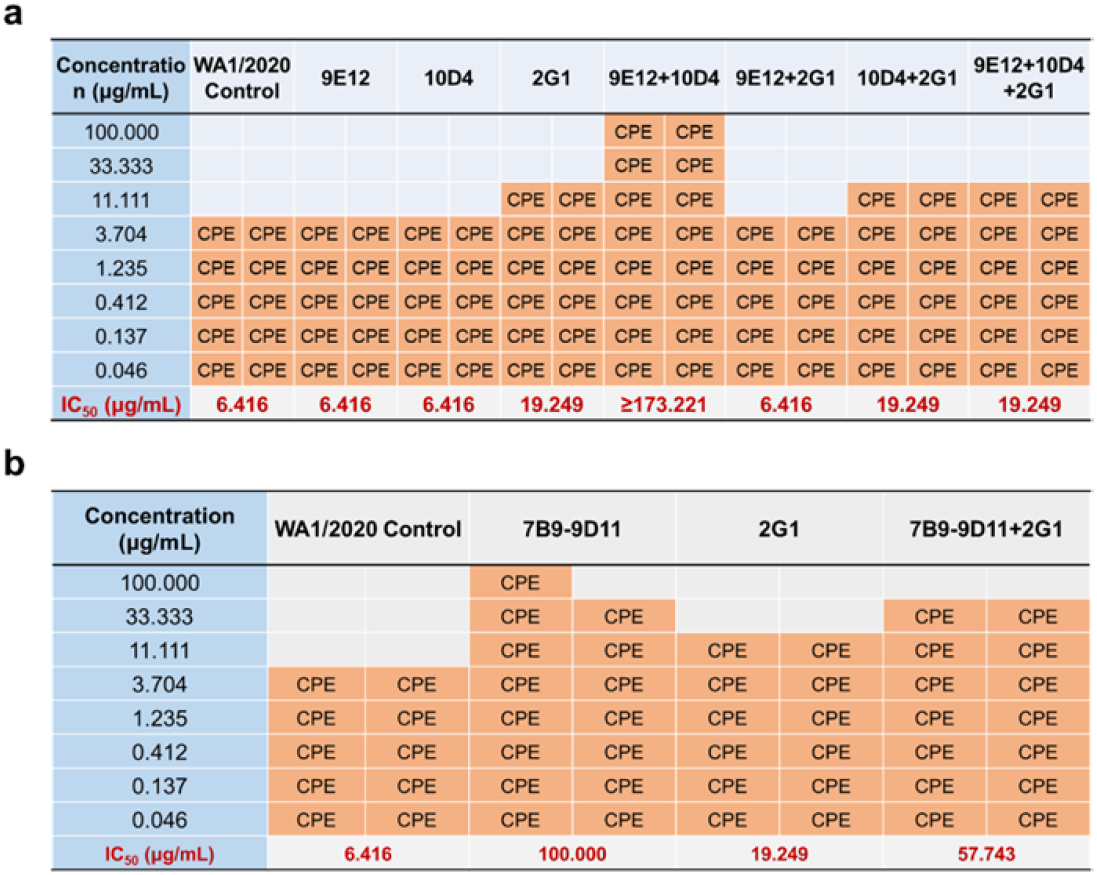
Comparison of ACE2 susceptibility of VSV-SARS-CoV-2 viruses that broke through the protection of antibody cocktails. After the viruses broke through the protection of 9E12+10D4+2G1 (**a**) and 7B9-9D11+2G1 (**b**), or reached the designed endpoint, the multi-passaged viruses were mixed with gradient diluted recombinant ACE2 protein and were used to infect Vero E6 cells. The cytopathic effect was observed and the antigenic affinity shift of the viral S protein were assessed according to the neutralizing alteration to recombinant ACE2.

### Deep sequencing identifying key viral escape sites

To understand how these two cocktails protect cells from mutant virus infection, we performed next generation sequencing to identify mutation sites in the S protein of the VSV-SARS-CoV-2 variants. Sole pressurized passaging by 9E12 generated three high proportion mutations, i.e., G485R, V503E and V615M (Figure 4a). While both antibodies 10D4 and 2G1 generated only one mutation site, K529N and F486V, respectively. The more mutations the virus generates may mean that the antibody is more difficult to be broken through, which could explain why the virus required more passages to break through the protection of 9E12 than 10D4 or 2G1 (Figure 2b). In the antibody cocktail by 9E12+10D4, the virus generated six mutation sites in S protein and two of them were in RBD region. Correspondingly, the virus did not break through 9E12+10D4 until the 29^th^ passage (Figure 2b), suggesting that 9E12+10D4 might be a relative tolerant combination that is difficult to be broken through. After sequencing the virus in the 2 μg/mL wells of 9E12+10D4+2G1, it was found that the virus emerged three mutations in S protein, i.e., H69R, G485R, and R683G. Although the three mutations did not completely break through the protection of 9E12+10D4+2G1, it weakened the neutralizing efficacy moderately. In terms of the 7B9-9D11+2G1 cocktail, four and five mutation sites came about from the 7B9-9D11 and 7B9-9D11+2G1 viruses respectively to weaken the antibody activity (Figure 4b).

**Figure 4.**
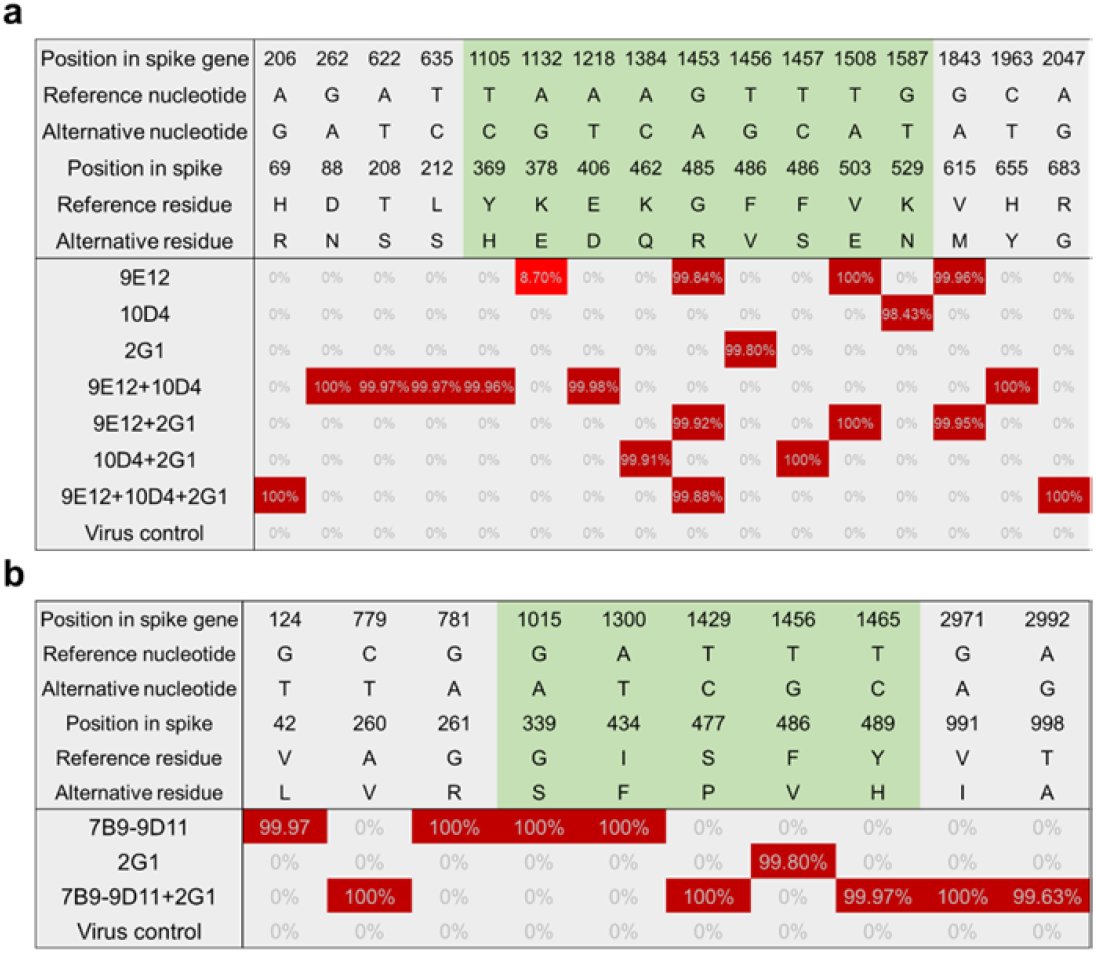
Deep sequencing of VSV-SARS-CoV-2 identifies key escape mutational sites for viruses pressurized replication from the 9E12+10D4+2G1 (a) and 7B9-9D11+2G1(b) cocktails. Medium containing pressurized viruses was collected and the next generation sequencing was performed to identify nucleotide mutations in the spike gene. The single nucleotide polymorphism (SNP) was analyzed with sufficient sequencing depth coverage to ensure the reliability. The RBD region is colored in light green and mutation sites are colored in red.

It is worth noting that the mutation sites of the viruses resistant to cocktails were not the addition of mutations from that of single antibodies. This may imply that more than one scheme for the virus to escape and involve in the overall conformational change of the S protein. Besides, most of these mutations are not found in VOC strains. This suggested that even if an antibody neutralizes most of the current variants, there may be still unknown mutations that can escape. Therefore, the anti-mutational potential of antibody combinations designed based on the current variants still needs to be explored reliably.

All these antibodies in our repertoire recognize the RBD region; however, there occurred non-RBD mutations. Studies suggested that mutations in the non-epitope region likely represent tissue culture adaptations^24,25^. We also infer that these mutations may together change the S protein in conformation and thus reduce antibody binding.

### Verifying the breadth of cocktails picked from VSV-SARS-CoV-2 passaging

Using a replication incompetent pseudovirus neutralizing system, we tested the activity of both 9E12+10D4+2G1 and 7B9-9D11+2G1 cocktails against all current VOC and VOI strains, as well as the SARS-CoV-1. As for monoclonal antibodies, 9E12 showed good neutralization against variants except for Omicron (BA.1); 10D4 inhibited the infection of most variants, especially the Omicron (half maximal inhibitory concentration IC_50_ = 1.4180 μg/mL), but its neutralization efficiency was relatively low, the IC_50_ values were generally over 0.1 μg/mL; 2G1 ultra-potently neutralized most variants, but had minimized activity against Omicron and had no activity against SARS-CoV-1 (Figure 5a). By contrast, the combinations of 9E12+10D4 and 10D4+2G1 not only neutralized Omicron, but also improved the efficiency against other variants. The 9E12+2G1 presented both high efficiency and the ability to neutralize SARS-CoV-1. Furthermore, the combination by 9E12+10D4+2G1 showed good neutralization to all variants, which was not achieved by monoclonal antibodies. Similar results were also observed on the 7B9-9D11+2G1 cocktail (Figure 5b). Antibody 2G1 increased the anti-viral efficiency and bispecific antibody 7B9-9D11 improved the anti-viral breadth.

**Figure 5.**
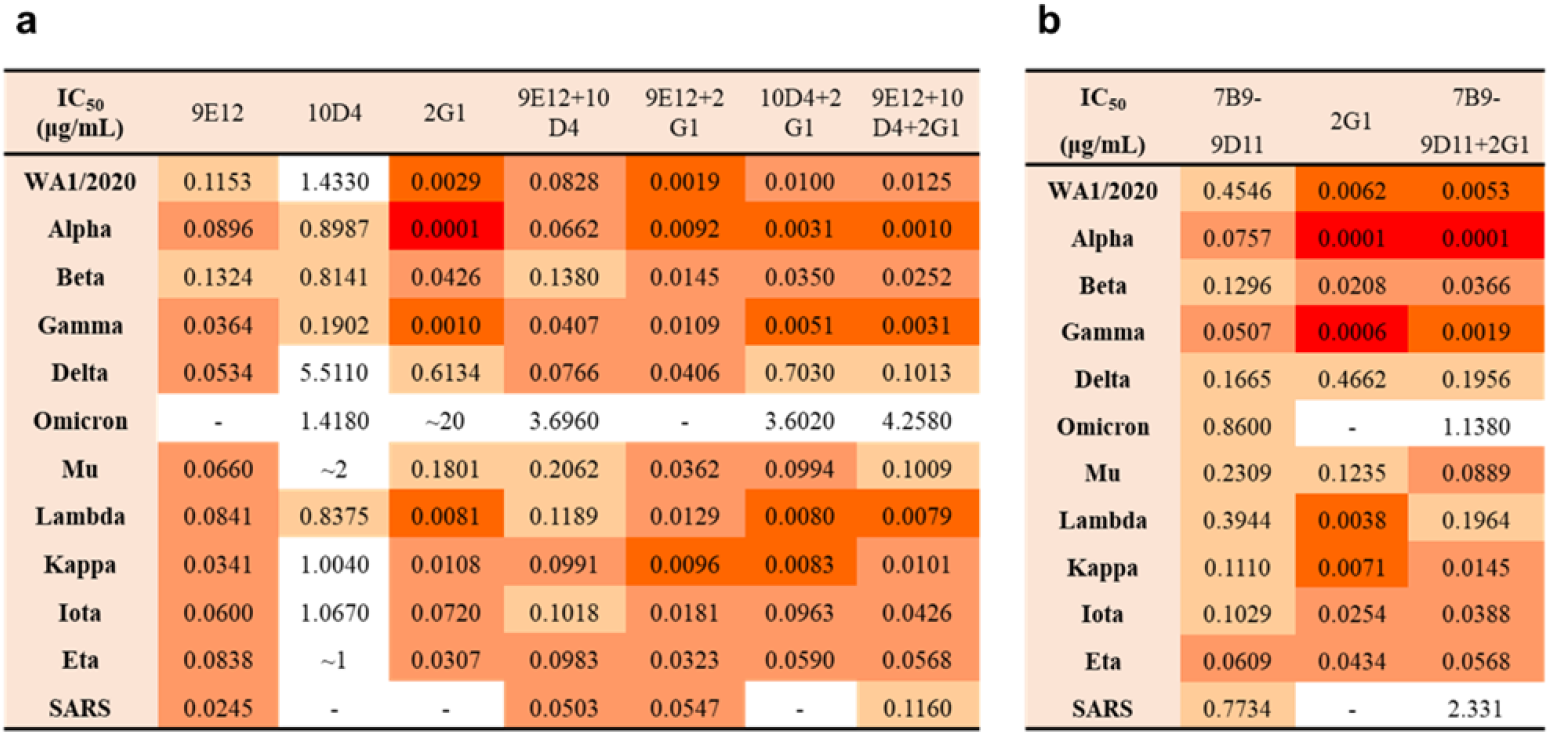
Evaluation of broad neutralizing ability of the 9E12+10D4+2G1 (a) and 7B9-9D11+2G1(b) cocktails in pseudo-typed SARS-CoV-2 and SARS-CoV-1 virus models. Antibodies were 10-fold serially diluted from 10^1^ μg/mL to 10^−5^ μg/mL and incubated with pseudo-typed viruses for 30 min, and then the mixture was added into a plate seeded with 293T-ACE2 cells. The infection of viruses to cells was quantified 48 hours later.

## Discussion

With the continuous emergency of SARS-CoV-2 variants, there are more and more mutations occured in the S region. The latest emerged variant Omicron sublineages have at least 30 substitutions in S region, which remarkably remodeled the S conformation^16^. The cryo-electron microscope structure revealed that mutations in Omicron enhanced the interaction between adjacent “up” and “down” conformation RBDs, which makes the S protein more stable and less conformational diversity^17^. Besides, mutations in Omicron also increased affinity with ACE2 and facilitated evasion from immunotherapy by neutralizing antibodies^18,26^. In fact, immune escape has been reported not only for Omicron but also for all VOC strains^27–30^. Antibody cocktails with breadth can effectively solve this issue. Several studies evaluated the antibody breadth using the VSV-SARS-CoV-2 system^31,32^; however, inadequate round of passaging is difficult to determine the tolerance of antibody cocktails accurately. We designed four cocktails, including monoclonal antibodies and bispecific antibodies, and explored their antiviral tolerance in the VSV-SARS-CoV-2 system. The experimental duration was more than 60 days thus the virus was allowed to produce enough mutations. We found that cocktails by 9E12+10D4+2G1 and 7B9-9D11+2G1 were able to protect Vero E6 cells from infection even experienced 30 passages. Although the virus reduced neutralizing activities by generating mutations, both ACE2 sensitivity and replication rate of the mutant virus decreased correspondingly, suggesting that these mutations may not become epidemic strains.

Based on our deep sequencing results of the S protein of VSV-SARS-CoV-2, we found that more mutations the virus needs to generate to break through the protection of more tolerated antibodies. The virus produced four mutations to break through the protection of 9E12, while only one mutation to break through 10D4 or 2G1. This was positively correlated with the rounds of passaging for their breakthrough. RBD antibodies can be classified into four types according to their germlines and antigen-binding models^7^. A number of antibodies in class 3 and class 4 with binding sites outside the RBD-ACE2 interface showed broad neutralization. Novel epitope antibodies targeting the back of the RBD-ACE2 interface identified recently and showed Omicron variant neutralization, which were called class 5 antibodes^16^. However, the breadth of a single antibody is always limited. In our study, although the virus generated mutations in the 9E12+10D4+2G1 and 7B9-9D11+2G1 cocktail samples, the two cocktails showed good neutralization from beginning to end. This result solidified the use of antibody cocktails to achieve a broad-spectrum anti-SARS-CoV-2 purpose.

We checked the neutralizing activity of these antibody candidates to SARS-CoV-1 but didn’t know to other variants in designing antibody cocktails. Through the pressurized culture against VSV-SARS-CoV-2, we obtained the 9E12+10D4+2G1 and 7B9-9D11+2G1 cocktails with good tolerance. To our surprise, the two cocktails neutralized all current VOC and VOI strains, as well as SARS-CoV-1 in the pseudovirus system. Which validated the reliability of using our VSV-SARS-CoV-2 system for screening and exploring antibody cocktails with breadth. Taken together, our study provided an approach to develop SARS-CoV-2 antibody cocktails with breadth and the feasibility of reasonably design of antibody cocktails to increase the anti-viral tolerance, which may work up the clinical solutions for tackling SARS-CoV-2 evasion.

## Materials and Methods

### Monoclonal antibody preparation

Plasmids pcDNA3.4 comprising heavy chains and light chains of antibodies were transiently expressed in ExpiCHO-S cells using the ExpiCHO™ expression system (Thermo Fisher). Briefly, the cells were expanded to 6 × 10^6^/mL the day before transfection and were then diluted to 3 × 10^6^/mL by ExpiCHO™ expression medium (Gibco). Plasmids were diluted to 20 μg/mL by cold OptiPRO medium (Gibco) and transfected into the ExpiCHO-S cells 17 h later according to the manufacturer’s instructions. The transfected cells were placed in a 37°C, 8% CO_2_ shaker, followed by Max Titer feeding 18 h later. Then, the cells were transferred to a 32 °C, 5% CO_2_ atmosphere. On day 5, the second Max Titer feeding was performed. When the cell viability reduced to about 90%, the medium was collected and centrifuged at 300 g for 10 min. The supernatant was subsequently further centrifuged at 7,000 g for 30 min to remove the cell debris. After filtered through 0.22 μm membranes (Millipore), the medium that containing antibodies was submitted to MabSelect Sure affinity purification. The full-length IgG antibodies were eluted with 100 mM citric acid (pH 3.0). Purified antibodies were stored in a cocktailed buffer consisting of 10 mM Histidine-HCl, 9% trehalose, and 0.01% polysorbate 80.

### Epitope binning

Competition enzyme-linked immunosorbent assay (ELISA) was performed to explore the epitope correlation of two antibodies. Briefly, the first antibody at the concentration of 2 μg/mL was coated on plates (BEAVER) and incubated at 4 °C overnight. Then, excess antibodies were washed away by PBS and blocked by 3% skim milk. SARS-CoV-2 S1 protein (Sino Biological) was biotinylated using an EZ-Link™ Sulfo-NHS-LC-LC-Biotin kit (ThermoFisher), followed by mixing with 50 μg/mL of the second competition antibodies or PBS blank control. After incubation at 37 °C for 1 h, plates were washed three times with PBS and the diluted Ultrasensitive Streptavidin-Peroxidase Polymer (Sigma) was added (1:2000) subsequently. Then the plates was incubated at 37 °C for 1 h again. TMB Single-Component Substrate Solution (Solarbio) was used to detect the S1 binding with the coated first antibodies. The absorbance at 450 nm was measured in an Infinite M200 PRO Multimode Microplate Reader (TECAN). Competitive percentage of two antibodies was calculated with reference to the PBS blank control.

### Bispecific antibody design and preparation

Based on our Bispecific Antibody by Protein Trans-splicing (BAPTS) platform, fragment A and fragment B of from two antibodies with poor epitope clash were chose to generate bispecific antibodies as previous described^22^. In brief, three plasmids pCDNA3.4 carrying full-length of heavy chain A, light chain A, and Fc region of heavy chain B jointing C-terminal of half intein were co-transfected into ExpiCHO-S cells to express fragment A; two plasmids pCDNA3.4 carrying light chain B and Fab region of heavy chain B jointing N-terminal of half intein were co-transfected to express fragment B. Fragment A and B were purified using HiTrap KappaSelect or HiTrap LambdaSelect (Cytiva). Glycine (100 mM, pH2.5) or acetic acid (100 mM, pH3.0) was used as elution buffer. For generating bispecific antibodies, fragment A and fragment B were mix in a mole ratio of 1:1.6. The reaction was happened in the presence of 2 mM Tris-(2-carboxyethyl)-phosphine and terminated by 2 mM dehydroascorbic acid after reaction for 4h at room temperature. The formed BsAbs were preliminary processed by Protein A resin (GE Healthcare) and subsequently purified employing HiTrap Capto MMC ImpRes (Cytiva). A linear elution scheme was tailored for Capto MMC purification, from 10 mM phosphate +10 mM Tris+50 mM NaCl (pH6.0) to 10 mM phosphate +10 mM Tris (pH9.0). The obtained BsAbs with purity >98% which was confirmed by size exclusion chromatography were concentrated and stored in −80°C.

### Antigen-binding and ACE2 competition ELISA

For quantifying the binding ability of neutralizing antibodies to SARS-CoV-1, the SARS-CoV-1 S protein (SinoBiological) was diluted to 1.0 μg/mL with ELISA Coating Buffer (Solarbio) and added to 96-well ELISA plates with 100 μL per well, and then placed in a 4 °C condition for overnight incubation. The next day, the plates were washed 4 times with 200 μL of PBST each time, followed by blocking with 200 μL of 3% skim milk. Neutralizing antibodies were diluted to 0.1 μg/mL in 1% bull serum albumin (Sigma) solution and 100 μL was added to each well. Subsequently, plates were incubated at 37 °C for 1 h. After further wash, 100 μL of goat anti-human IgG (Fc specific)-Peroxidase antibody (1 : 5000 dilution, Sigma) was added and incubated for another 1 h at 37 °C. Then 100 μL of TMB substrate (Solarbio) was added and incubated in the dark for 15 min to visualize the binding of the antibody to the SARS-CoV-1 antigen. The chromogenic reaction was terminated by adding 50 μL of stop buffer (Solarbio) and the absorbance at 450 nm was immediately read in the microplate reader (TECAN).

For investigating competitive binding of antibodies in the cocktail to trimeric SARS-CoV-2 S protein, recombinant hACE2-Fc protein was biotinylated using the EZ-Link sulfo-nhs-biotin kit (ThermoFisher) according to the manufacturer’s instructions. ELISA plates were coated with SARS-CoV-2 S protein in the same way as described above. The work concentration of biotinylated ACE2 was optimized, and the 80% of maximal effect (EC_80_) concentration of antigen binding was used for this experiment. Equal volumes of antibodies and biotinylated ACE2 (50 μL : 50 μL) were mixed and added into the plates. Detection was performed with Ultrasensitive Streptavidin-Peroxidase Polymer (1 : 2000, Sigma). PBS and non-biotinylated ACE2 protein substituted for competing antibodies were served as blank control and positive control, respectively.

### Surface plasmon resonance (SPR)

The BIAcore 8K system was used to detect the binding of antibodies to the S trimer of SARS-CoV-2 WA1/2020 (AcroBiosystems). The S protein trimer was covalently coupled to the CM5 sensor chip using an amino coupling kit (Cytiva). After the response value reached about 70 relative unit (RU), the coupling was stopped and the excess S protein was washed away, followed by blocking unbound sites with ethanolamine (Cytiva). Antibodies were diluted to 50 μg/mL with HBS-EP buffer (Cytiva). In order to detect whether more than one antibodies can bind to S protein simultaneously, the first antibody was loaded into the system and flowed for 600 s to meet saturated binding. Then the second antibody was loaded into the identical channel and flowed for another 600 s. If there was a third or more antibody, it was proceeded in the same way. Channels flowing with the same antibody as previous rounds were used as control. The simultaneous binding of antibodies was judged by the accumulation of response values from different rounds of flow.

### VSV passaging

The replication competent vesicular stomatitis virus incorporating the spike of SARS-CoV-2 (VSV-SARS-CoV-2) was generated as described previously^33^, under the technical guidance of Professor Tao Sun of Shanghai Jiao Tong University. For testing the breadth of antibody cocktails, Vero E6 cells were seeded into 96-well plates with 2.5 × 10^4^ cells per well. The next day, antibodies were 5-fold serially diluted starting from 100 μg/mL in DMEM medium supplementing 10% FBS (Gibco). Wells without antibodies or containing the same concentrations of REGN10933 were set as control. Then 50 μL of the antibody dilution was mixed with 50 μL of VSV-SARS-CoV-2 in the final multiplicity of infection (MOI) of 0.01. After incubated at room temperature for 30 min, the virus-antibody mixture was added to the cells and grown for two days at 37°C, 5% CO_2_. The cytopathic effect (CPE) of cells was examined under a microscope (Nikon). If the CPE in a well reached 20%, the virus was considered to have broken through the protection of antibodies. The medium supernatant of the wells with the highest antibody concentration that has been broken was collected and was diluted 12 times for the next round of passaging. After the virus broke through the highest antibody concentration wells or reached the designed endpoint, the medium supernatant was collected for next generation sequencing. Single nucleotide polymorphisms (SNP) of the S region was analyzed according to the sequencing results.

### VSV characterization

The replication rates of mutant viruses were measured according to the increase in number of viral particles in one generation. In 24-well plates, 1 × 10^5^ Vero E6 cells were seeded per well and cultured for 24 h. The next day, each mutant VSV-SARS-CoV-2 was added to the cells with duplicates at MOI = 0.01 per well. Then the plates were gently tapped to mix the virus evenly, followed by incubation for another 12 h. Cells were disassociated from the plate using 100 μL of trypsin (Gibco) and fixed by 4% paraformaldehyde (Solarbio) at the time points of 0 h, 1 h, 2 h, 3 h, 4 h and 5 h. The virus particles that have infected cells were quantified by analyzing the green fluorescent protein reporter that was comprised into the viral genome in a flow cytometer. The replication rate of the virus was calculated based on the mean fluorescence intensity.

For investigating the ACE2 susceptibility of mutant VSV-SARS-CoV-2, Vero E6 cells were seeded into 96-well plates at 2.5 × 10^4^ cells per well. The next day, the recombinant ACE2-Fc protein which was obtained from ExpiCHO™ expression system (Thermo Fisher) was 5-fold serially diluted in DMEM medium supplementing 10% FBS (Gibco). The VSV-SARS-CoV-2 viruses were diluted by medium and mixed with equal volume of ACE2-Fc dilutions. The final MOI was 0.01. Then, 100 μL of the mixture was added to the cells, followed by further incubation at 37 °C under 5% CO_2_ condition for 48 h. Under a microscope (Nikon), the CPE of the cells was checked and the IC_50_ values were calculated according to the equation of IC_50_ = Antilog (D - C × (50 - B) / (A - B)). Where A means percent inhibition higher than 50%, B represents percent inhibition less than 50%, C is log_10_ (dilution factor), and D is log_10_ (sample concentration) that the inhibition is less than 50%.

### Pseudovirus neutralization

The pseudovirus neutralization study was performed as described previously^21^. In brief, 293T-ACE2 cells were plated at a density of 1 × 10^4^ cells per well into a 96-well white transparent bottom plate, and cultured overnight at 37 °C in 5% CO_2_. Serial 10-fold dilutions of neutralizing antibodies were mixed with pseudoviruses. After incubation at 37 °C for 30 min, the mixture was added into the 293T-ACE2 cells. The medium was replaced with fresh medium 6 h later. All operations involving pseudoviruses were performed in a biosafety level 2 laboratory in the School of Pharmacy, Shanghai Jiao Tong University. After further culturing for 48 hours, 50 μL of ONE-Glo luciferase substrate (Promega) was added to each well, and the fluorescence intensity was detected on a microplate reader (TECAN) immediately. Data were analyzed using GraphPad Prism software and the neutralization efficiencies of antibodies were calculated according to the four-parameter nonlinear fitting results.

## Acknowledgments

Authors would like to acknowledge following organizations and individuals for their assistances in the preparation of the manuscript: Professor Tao Sun of SJTU for providing technical and material supports in generating the VSV-SARS-CoV-2 system. This work was funded by the National Natural Science Foundation of China (81773621, 82073751 to J.Z.); the National Science and Technology Major Project “Key New Drug Creation and Manufacturing Program” of China (No.2019ZX09732001-019 to J.Z.); the Key R&D Supporting Program (Special support for developing medicine for infectious diseases) from the Administration of Chinese and Singapore Tianjin Eco-city to Jecho Biopharmaceuticals Ltd. Co.; Zhejiang University special COVID-19 grant 2020XGZX099 and Shanghai Jiao Tong University “Crossing Medical and Engineering” grant 20X190020003 to JZ.

